# Difficulties with speech-in-noise perception related to fundamental grouping processes in auditory cortex

**DOI:** 10.1101/814913

**Authors:** Emma Holmes, Peter Zeidman, Karl J. Friston, Timothy D. Griffiths

## Abstract

In our everyday lives, we are often required to follow a conversation when background noise is present (“speech-in-noise” perception). Speech-in-noise perception varies widely—and people who are worse at speech-in-noise perception are also worse at fundamental auditory grouping, as assessed by figure-ground tasks. Here, we examined the cortical processes that link difficulties with speech-in-noise perception to difficulties with figure-ground perception using functional magnetic resonance imaging (fMRI). We found strong evidence that the earliest stages of the auditory cortical hierarchy (left core and belt areas) are similarly disinhibited when speech-in-noise and figure-ground tasks are more difficult (i.e., at target-to-masker ratios corresponding to 60% than 90% thresholds)—consistent with increased cortical gain at lower levels of the auditory hierarchy. Overall, our results reveal a common neural substrate for these basic (figure-ground) and naturally relevant (speech-in-noise) tasks—which provides a common computational basis for the link between speech-in-noise perception and fundamental auditory grouping.

## Introduction

One of the greatest challenges of everyday listening is the requirement to understand speech when background noise is present (“speech-in-noise” perception). People vary widely in their ability to understand speech-in-noise, yet we do not fully understand the processes that contribute to this variability. A widely used clinical measure of peripheral hearing ability—the pure-tone audiogram—is unable to fully account for speech-in-noise difficulties (Cooper and Gates, 1991; Hind et al., 2011; Holmes and Griffiths, 2019; Kumar et al., 2007). Yet, recent evidence suggests that individual variability in central processes relates to speech-in-noise perception (Holmes and Griffiths, 2019). Here, we use functional magnetic resonance imaging (fMRI) to examine the cortical processes that relate to speech-in-noise difficulty.

Holmes and Griffiths (2019) demonstrated that central auditory grouping processes also vary widely between people and can help to explain individual differences in speech-in-noise perception. They assessed fundamental grouping processes using stochastic figure-ground perception—which tests the ability to track pure tones that retain the same frequency over time (the ‘figure’) amongst a ‘background’ of tones of random frequencies (Teki et al., 2013, 2011). Holmes and Griffiths (2019) varied the target-to-masker ratio between figure and background tones to estimate thresholds for discriminating gaps that occurred in the figure or background components. They found that figure-ground thresholds helped to explain individual differences in speech-in-noise perception, after accounting for differences in peripheral hearing predicted by the pure-tone audiogram. In other words, people who are worse at speech-in-noise perception are worse at figure-ground perception.

Figure-ground perception is non-linguistic—and the link between figure-ground and speech-in-noise perception likely reflects a shared reliance on fundamental grouping processes, which are used to segregate figure from background tones and to segregate target speech from other sounds (including competing speech). Similar to speech-in-noise perception, the ‘figure’ and ‘background’ tones used by Holmes and Griffiths (2019) overlap in frequency, so the figures cannot be detected based on simple spectral separation. Instead, proposed figure-ground mechanisms are based on binding figure elements of different frequencies—based on the detection of temporal coherence between figure elements, as measured by cross correlation (Shamma et al., 2011; Teki et al., 2013). Auditory cortex contains neurons that respond to multiple frequencies (Elhilali et al., 2009a; Rauschecker, 1998; Wang, 2013), and thus could plausibly represent figures containing multiple frequency elements. However, any account of the brain basis for the behavioural correlation between figure-ground and speech-in-noise perception must also account for the fact that these are both active tasks, and the task may interact with the stimulus representation.

In studies in which there is no task or an irrelevant task, processing of figure-ground stimuli is associated with activity in high-level auditory-cortex and the intraparietal sulcus. Teki et al. (2011) showed with fMRI that figure properties associated with increased salience (i.e., longer figures or figures with more frequency components), produce greater activity in the superior temporal sulcus (STS) and inferior parietal sulcus (IPS), without a task. Additionally, longer figures produced more activity in auditory cortex: in the right planum temporale (PT). When Teki et al. (2011) specifically examined early auditory cortex (human homologues of ‘core’ designated Te 1.0 and ‘belt’ designated Te 1.1 and Te1.2; Morosan et al., 2001), there was no evidence for modulations of activity by these figure manipulations. A magnetoencephalography (MEG) study that used an irrelevant task (Teki et al., 2016) similarly demonstrated that activity of high-level auditory cortex and IPS were modulated by the same figure manipulations that increase salience. Figure detection in macaque monkeys has also been localised to higher parabelt areas of auditory cortex (Schneider et al., 2018). These studies are consistent with stimulus-driven effects in auditory cortex, although additional attentional effects on the brain signal evoked by figure-ground stimuli have been demonstrated using EEG (O’Sullivan et al., 2015). The brain locus for attentional effects has also been assessed using MEG (Molloy et al., 2019), which revealed an effect of attentional load on activity in early auditory cortex. In the present fMRI study, we were specifically interested in task effects in early auditory cortex that might be relevant to both figure-ground analysis and speech-in–noise perception. To this aim, we compared brain activity (here, estimated using blood-oxygen level-dependent [BOLD] activity) during active figure-ground analysis and speech-in-noise perception, and modelled responses in the auditory cortical hierarchy to establish the mechanism by which task effects operate.

Previous studies of speech-in-noise perception have revealed responses in a wide variety of areas, including parts of auditory cortex that have been associated with figure-ground perception. When more difficult listening conditions (e.g., speech at a lower target-to-masker ratio [TMR]) are compared with easier listening conditions (e.g., at a higher TMR or without background noise), the posterior STS (Eckert et al., 2016) and bilateral superior temporal gyrus (STG) (Wong et al., 2008) show greater activity. Although primary (core) auditory cortex has typically been absent in studies comparing different speech-in-noise conditions, it shows a relationship with the intelligibility of speech presented alone: it shows greater activity for clear relative to degraded (vocoded) speech, greater activity for degraded than unintelligible speech, and greater activity when the intelligibility of degraded speech is enhanced by presenting a matching word prime (Wild et al., 2012). Putative belt areas of auditory cortex show similar patterns: Anterolateral Heschl’s gyrus shows a preference for clear compared to vocoded speech (Nourski et al., 2019), and posterior Heschl’s gyrus shows a relationship with speech intelligibility in signal-correlated noise (Davis et al., 2011). Beyond auditory cortex, speech-in-noise tasks engage fronto-parietal areas (Eckert et al., 2016; Hill and Miller, 2010) that are commonly associated with attention—in particular, the inferior frontal gyrus, but also the inferior frontal sulcus and middle frontal gyrus (Binder et al., 2004; Davis et al., 2011; Scott et al., 2004; Zekveld et al., 2006). Activity has also been observed in cingulo-opercular regions, including the insula, frontal operculum, and cingulate gyrus (Eckert et al., 2016; Vaden Jr et al., 2016, 2013).

Based on these studies, we anticipated that the greatest overlap of functional integration between figure-ground and speech-in-noise perception would occur in auditory cortex. We used fMRI to measure BOLD activity in combination with dynamic causal modelling (DCM) to examine the common (i.e., shared) changes in effective connectivity engaged by active speech-in-noise and figure-ground tasks. Regarding BOLD effects, we predicted the two tasks would activate distinct higher-level regions beyond auditory cortex (e.g., IPS, IFG, and cingulo-opercular regions), but we predicted overlapping activations in early auditory cortex. To ensure that behavioural performance did not explain differences between the two tasks, we selected target-to-masker ratios for which accuracy was matched. To assess the functional anatomy of the shared behavioural relationship, we selected two different target-to-masker ratios—which, here, correspond to the salience of the figure and speech or, in other words, the difficulty of the tasks—that led to different performance levels. For the DCM analysis, we first assessed top-down connectivity from higher to lower levels of the auditory cortical hierarchy when speech-in-noise and figure-ground perception are more difficult. This would be consistent with the longstanding idea that, when speech is perceptually ambiguous due to noise, listeners rely on (higher-level) prior expectations to a greater extent (e.g., Marslen-Wilson and Tyler, 1980; Mattys et al., 2012; McClelland and Elman, 1986; Norris et al., 2015; Norris and Mcqueen, 2008; Peelle, 2017). We were also interested to quantify changes in effective connectivity within early sub-regions of auditory cortex, which might explain attentional and task effects suggested by earlier studies. An advantage of using DCM is that it allows us to compare—using Bayesian model comparison—models that do and do not contain task-specific effects of difficulty. If models without task-specific effects have greater evidence, we can conclude that greater difficulty in both tasks are mediated by similar changes in directed neuronal coupling.

## Results

Behaviour (d′) in the scanner followed the expected patterns (see Figure 1). A two-way within-subjects ANOVA showed a significant main effect of Difficulty [F(1, 43) = 19.40, *p* < .001, *ω*_*p*_^2^ = .29] and no significant main effect of Task [F(1, 43) = .56, *p* = .46, *ω*_*p*_^2^ = -.01]. The interaction between Task and Difficulty was not significant either [F(1, 43) = .95, *p* = .34, *ω*_*p*_^2^ < .01]. These results confirm that the difficulty manipulation was successful, and show that performance did not differ significantly between the two tasks.

**Figure 1.**
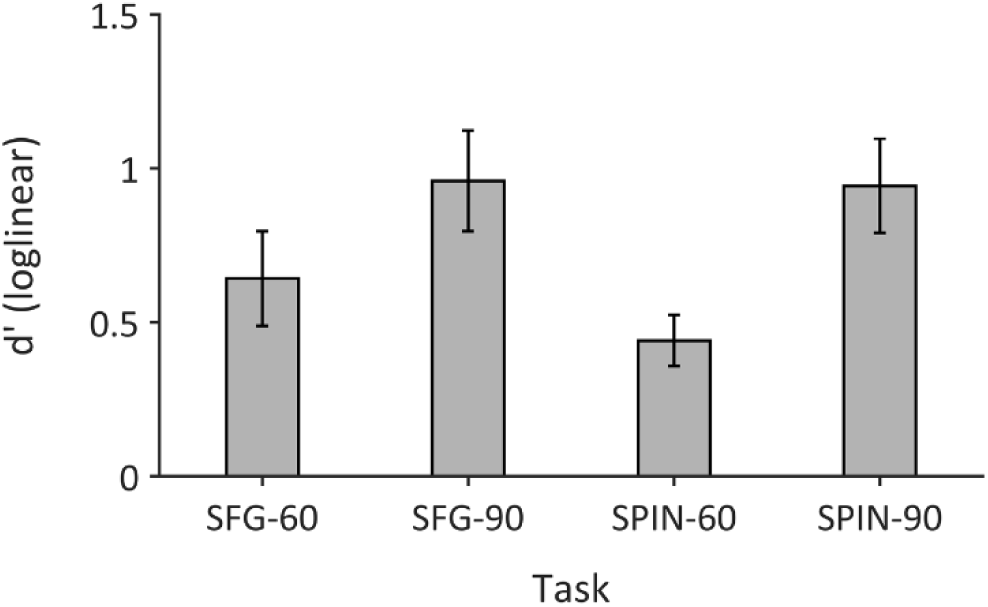
Sensitivity in the stochastic figure-ground (SFG) and speech-in-noise (SPIN) tasks, at the two TMRs corresponding to 60% and 90% thresholds. Error bars display ±1 standard error of the mean.

### Figure-ground perception predominantly activates a sub-set of the areas for speech-in-noise perception

The univariate analyses showed significant differences in activity between the two tasks. Table 1 lists the statistics and Figure 2A–B displays the thresholded SPMs. The speech-in-noise task was associated with greater activity in bilateral STG, the left precentral gyrus, and the right cerebellum than the figure-ground task; these results all survived a threshold of *p* < .001 after family-wise error (FWE) correction. Whereas, the opposite contrast yielded only two significant voxels, and they did not survive a *p*_*FWE*_ < .001 threshold. The finding that greater activity was revealed by the SPIN > SFG contrast than the SFG > SPIN contrast implies that the figure-ground task predominantly activates a sub-set of the areas that are active for the speech-in-noise task.

**Table 1.** Contrasts between the speech-in-noise (SPIN) and stochastic figure-ground (SFG) tasks. Statistical analyses were conducted at the group level using one-sample t-tests, and were thresholded at p = .05 after correcting for family-wise error (FWE). Peak locations were determined using Neuromophometrics (Neuromorphometrics, Inc.; http://neuromorphometrics.com/).

| Contrast | Peak location | X (mm) | Y (mm) | Z (mm) | *t* | *p*<sub>FWE</sub> |
| --- | --- | --- | --- | --- | --- | --- |
| SPIN > SFG | Left STG | -63 | -36 | 6 | 8.56 | < .001 |
|  |  | -60 | -3 | -3 | 8.00 | < .001 |
|  |  | -57 | -12 | 0 | 7.97 | < .001 |
|  | Left precentral gyrus | -51 | -12 | 42 | 7.95 | < .001 |
|  | Right STG | 60 | -21 | 0 | 7.48 | < .001 |
|  |  | 66 | -15 | 0 | 7.13 | < .001 |
|  |  | 63 | -6 | -6 | 6.97 | < .001 |
|  | Right cerebellum | 24 | -60 | -21 | 6.99 | < .001 |
|  | Right cerebellum | 15 | -66 | -12 | 5.91 | .011 |
|  | Left inferior temporal gyrus | -45 | -57 | -9 | 5.82 | .014 |
|  | Left putamen | -30 | -15 | -3 | 5.61 | .029 |
|  | Left middle frontal gyrus | -39 | 6 | 36 | 5.55 | .036 |
|  | Cerebellar Vermal Lobules I-V | 6 | -57 | -3 | 5.53 | .038 |
|  | Left STG | -51 | -9 | -12 | 5.52 | .040 |
|  | White matter | 9 | -75 | 24 | 5.47 | .047 |
| SFG > SPIN | White matter | -21 | -6 | 51 | 5.72 | .020 |
|  | Left superior parietal lobule | -36 | -39 | 42 | 5.46 | .048 |

**Figure 2.**
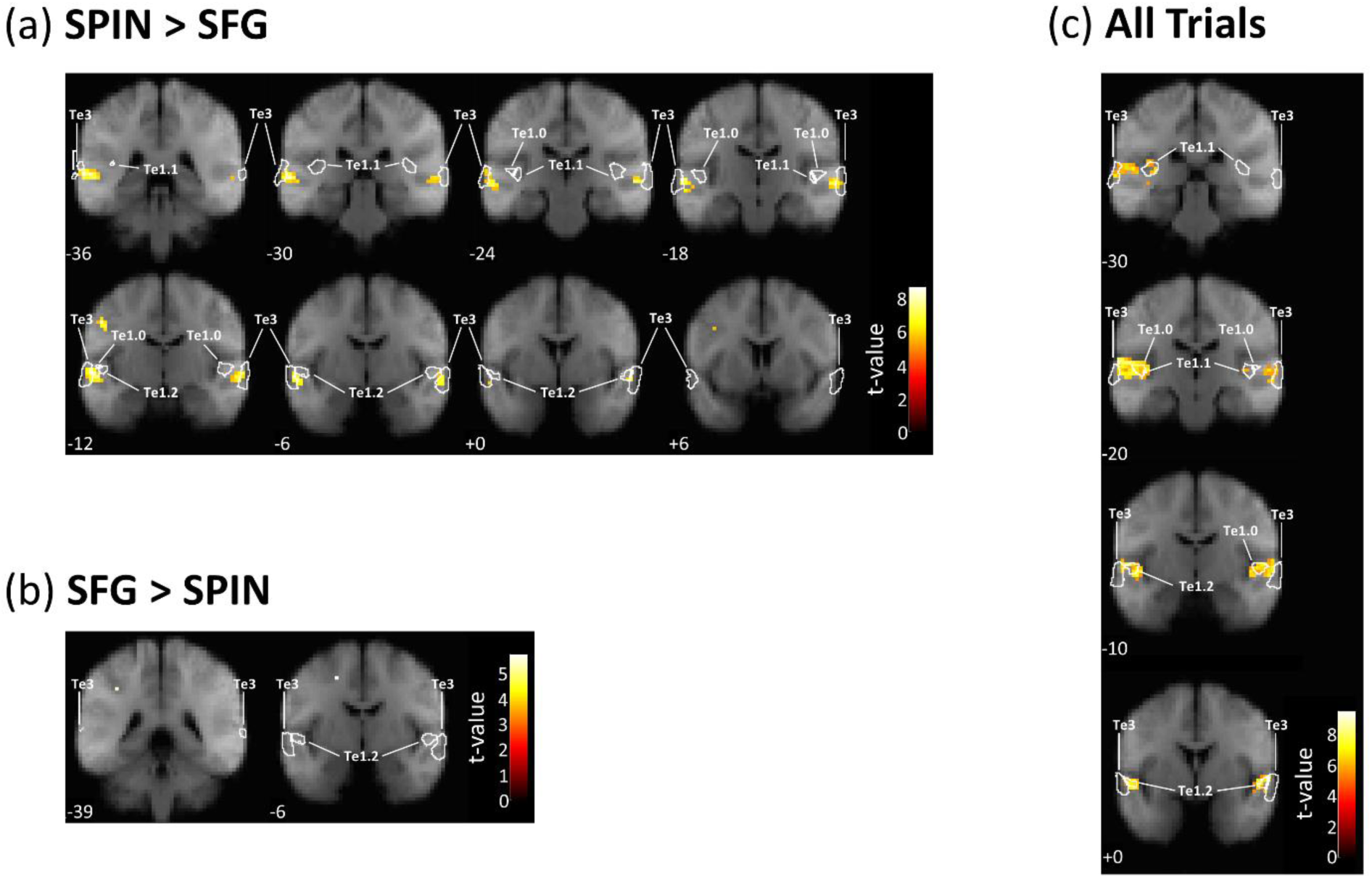
Group-level contrasts, thresholded at p < .05 (family-wise error corrected) and superimposed on coronal sections of the average (N = 44) T1-weighted structural image. (a) Voxels showing greater activity for the speech-in-noise (SPIN) than stochastic figure-ground (SFG) task. (b) Voxels showing greater activity for the stochastic figure-ground than speech-in-noise task. (c) Voxels for the All Trials > baseline contrast. MNI co-ordinates (in mm) are displayed below each coronal section. White lines show the outlines of anatomical maps corresponding to areas Te1.0, Te1.1, Te1.2, and Te3.

None of the other contrasts (main effect of Difficulty or Task–Difficulty interactions) revealed activity below the *p* = .05 FWE threshold.

### Difficulty with speech-in-noise and figure-ground perception lead to similar disinhibition in auditory cortex

To determine nodes for the DCM analysis, we contrasted All Trials against baseline. Table 2 lists the locations of voxels that survived the p < .05 FWE threshold, and Figure 2C shows the locations on four coronal slices. The voxels that survived a correction of p < .001 FWE were located on the superior temporal lobe—with peaks in the left transverse temporal gyrus (Heschl’s gyrus), left planum temporale, and bilateral planum polare. Thus, for the DCM analyses we focussed broadly on these parts of auditory cortex as areas of interest. We segmented the functional activity into anatomical regions of the auditory hierarchy: bilateral Te1.0, Te1.1, Te1.2, and Te3 (Morosan et al., 2005, 2001).

**Table 2.** Contrast of All Tasks > baseline. Statistical analyses were conducted at the group level using one-sample t-tests, and were thresholded at p = .05 after correcting for family-wise error (FWE). Peak locations were determined using Neuromophometrics (Neuromorphometrics, Inc.; http://neuromorphometrics.com/).

| Peak location | X (mm) | Y (mm) | Z (mm) | *t* | *p*<sub>FWE</sub> |
| --- | --- | --- | --- | --- | --- |
| Left planum temporale | -60 | -18 | 9 | 9.55 | < .001 |
| Left transverse temporal gyrus | -51 | -15 | 6 | 9.45 | < .001 |
| Left planum polare | -45 | -12 | 0 | 8.75 | < .001 |
| Right planum polare | 57 | 3 | 0 | 9.01 | < .001 |
| Right cerebral white matter | 63 | -12 | 3 | 8.89 | < .001 |
| Right planum polare | 51 | -3 | -3 | 7.91 | < .001 |
| Right anterior insula | 30 | 24 | 0 | 6.74 | .001 |
| Right frontal operculum | 33 | 18 | 9 | 5.60 | .030 |
| Left supplementary motor cortex | -6 | 18 | 45 | 6.31 | .003 |
| Left opercular IFG | -48 | 21 | 24 | 5.86 | .013 |
| Left superior parietal lobule | -27 | -54 | 48 | 5.60 | .031 |
| Left anterior insula | -39 | 18 | -3 | 5.58 | .032 |
| Left anterior insula | -33 | 21 | 0 | 5.58 | .032 |
| Right planum polare | 42 | -15 | -6 | 5.55 | .036 |
| Left cerebral white matter | -39 | -30 | 0 | 5.55 | .036 |
| Right cerebellum | 21 | -66 | -51 | 5.52 | .040 |
| Right opercular IFG | 48 | 9 | 21 | 5.48 | .045 |
| Right opercular IFG | 57 | 21 | 15 | 5.48 | .045 |

Table 3 displays the parameters of interest in the final group-level DCM, after Bayesian model reduction and averaging. These parameters quantified the modulatory effects of Difficulty and the interaction between Task and Difficulty on each connection. Only two of the parameters had high probabilities (Pp > .95). These two parameters correspond to the modulation of intrinsic (i.e., self) connections for left Te1.0 and left Te1.1 by the main effect of Difficulty (Figure 3). The values of these parameters were -.31 and -.30, respectively, which indicate a decrease in self-inhibition associated with the experimental effects.

**Table 3.** Parameters in the Dynamic Causal Model (DCM) after Bayesian model reduction and averaging. Target-to-masker ratios (TMR) in each of the four conditions (SPIN-90, SPIN-60, SFG-90, SFG-60) were included as regressors. The connection strengths and posterior probabilities associated with the parameters (based on the free energy with and without the parameter) are displayed in the final columns. For intrinsic parameters, the prior connection strength is -.5 Hz— reflecting an inhibitory self-connection; therefore, values smaller than -.5 reflect disinhibition relative to the prior. For extrinsic parameters, the prior is set to 0. Parameters with probabilities < .95 are listed in *italics*.

|  | Parameter | Connection strength (Hz) | Posterior probability |
| --- | --- | --- | --- |
| Commonalities | Difficulty modulation: Left Te1.0 (intrinsic) | $-.31$ | $>.99$ |
| | Difficulty modulation: Left Te1.1 (intrinsic) | $-.30$ | $>.99$ |
| | *Difficulty modulation: Left Te1.3 (intrinsic)* | $-.38$ | $.94$ |
| | *Difficulty modulation: Right Te1.1 (intrinsic)* | $-.42$ | $.71$ |
| SFG-60 TMR | *Interaction modulation: Left Te1.1-Left Te1.2* | $-.02$ | $.78$ |
| SPIN-60 TMR | *Difficulty modulation: Left Te1.2 (intrinsic)* | $-.53$ | $.76$ |
| | *Difficulty modulation: Right Te1.0 (intrinsic)* | $-.53$ | $.73$ |
| | *Difficulty modulation: Right Te1.2 (intrinsic)* | $-.53$ | $.81$ |

**Figure 3.**
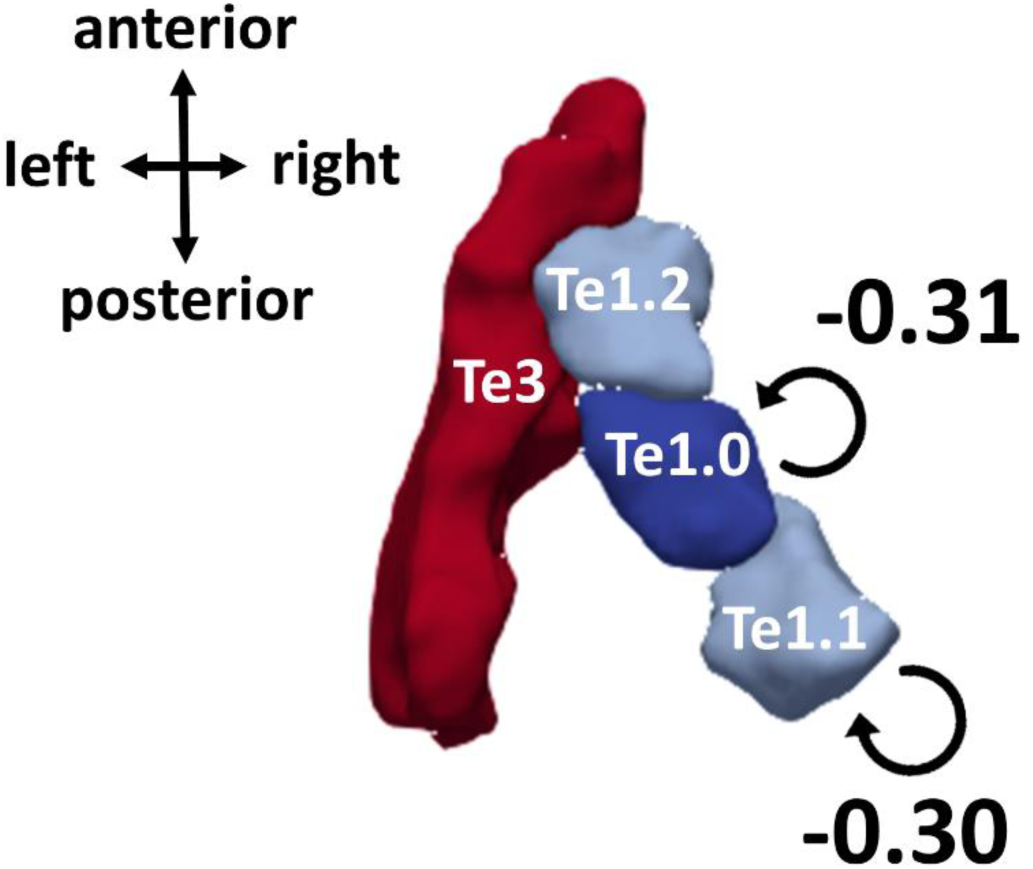
Group-level Dynamic Causal Model (DCM), after Bayesian model reduction and averaging. Parameters that survive a threshold of Posterior Probability (P_p_) > .95 are displayed: These correspond to modulations of intrinsic connectivity by the main effect of Difficulty in left Te1.0 and 1.1. Their connection strengths (in Hz) are displayed in the figure. Results are displayed on a 3D reconstructed surface of the anatomical regions of interest in the left hemisphere, which was generated using ITK-SNAP (www.itksnap.org; Yushkevich et al., 2006) and ParaView (www.paraview.org; Ahrens et al., 2005). Note that right-hemisphere homologues were included in the DCM, but none of the parameters had posterior probabilities greater than .95.

Interestingly, all of the parameters corresponding to modulations of connectivity by the Task– Difficulty interaction were pruned away, as were as all of the modulations of extrinsic connectivity— and therefore, these parameters are not present in the final model. In other words, the model evidence was greater when these connections were ‘switched off’ than when they were ‘switched on’. Thus, there is very strong evidence that, in auditory cortex, effects of Difficulty on effective connectivity are shared between the speech-in-noise and figure-ground tasks. The evidence does not support partially shared effects of Difficulty (which would be indicated by modulations of effective connectivity by both Difficulty and the Task-Difficulty interaction) or separate effects (which would be indicated by modulations of effective connectivity by the Task-Difficulty interaction only).

Some of the model parameters covaried with the TMR in one of the four conditions, although it is worth noting that their posterior probabilities were low (≤ .81), and their connection strengths differed minimally from the prior.

## Discussion

Our results demonstrate that speech-in-noise and figure-ground perception rely on similar cortical processes. Using DCM, we found that greater difficulty in both tasks leads to similar disinhibition at lower levels of the auditory cortical hierarchy: We found strong evidence (> 99% probability) that left Te1.0 and left Te1.1 are disinhibited when the tasks are more difficult—indicating that these regions likely increase their gain when speech or figures are difficult to follow. Importantly, we found the best model had no modulations by the Task–Difficulty interaction, suggesting that the best explanation of the data is that effective connectivity in auditory cortex is common to the two tasks, rather than task-specific. Overall, these results provide evidence for a common cortical substrate in early auditory cortex that could explain why people who find speech-in-noise perception difficult also find figure-ground perception difficult (Holmes and Griffiths, 2019).

The results showed that lower-level intrinsic connectivity, rather than top-down connectivity from higher to lower areas, was modulated by task difficulty—consistent with previous reports that activity in early auditory cortex is modulated by attentional load during figure-ground analysis (Molloy et al., 2019). Previous studies of visual scene analysis that used DCM similarly found that lower-level intrinsic connections, rather than top-down connections, offered the best explanation for modulations of BOLD activity by the ‘noisiness’ of a random dot motion stimulus (Adams et al., 2016, 2015). Intrinsic connections in DCM are rate constants, which control the rate of decay in a region, and disinhibitory modulations indicate slower decay. In other words, disinhibition indicates that an area becomes more excitable. In predictive coding formulations of perceptual synthesis, this is usually interpreted in terms of assigning more precision to prediction errors, so that they have greater influence on evidence accumulation or Bayesian belief updating. From a psychological perspective, this is usually thought of in terms of attentional selection (Auksztulewicz and Friston, 2015; Bauer et al., 2014; Brown et al., 2013; Kanai et al., 2015; Vossel et al., 2015, 2014).

Thus, our results imply that when figure-ground and speech-in-noise perception are more difficult— in other words, when figures and speech are less salient—we ‘listen harder’ or ‘pay more attention’ to lower levels of the auditory hierarchy. A likely mechanism for disinhibition of intrinsic connections is increased neuronal gain or excitability, which is well-established in auditory cortex (Rabinowitz et al., 2011). This could be realized by communication through coherence, by spiking inter-inhibitory neurons equipped with N-methyl-D-aspartate (NMDA) receptors, or by modulations through acetylcholine (ACh). Broadly speaking, these mechanisms are unlikely to be synapse-specific and would therefore more likely manifest in changes in intrinsic than extrinsic connectivity. Modulations of particular extrinsic (e.g., top-down) connections would be more consistent with synapse-specific effects.

Modulations of lower-level intrinsic connectivity are also consistent with the results of some previous studies examining speech perception. For example, Mattys et al. (2005) found that when speech is accompanied by white noise, lexical decisions are based on lower level (word stress) cues rather than higher level (e.g., context) cues. In addition, several studies have proposed that listening challenges during speech perception are realised by feedforward rather than feedback processes. For example, Davis et al. (2011) used fMRI to measure responses to spoken sentences. They found that listening challenges introduced by presenting speech at a low TMR or by presenting semantically ambiguous sentences were associated with early responses in anterior STG, which preceded later changes in higher areas. They interpret their results as reflecting a greater demand on internal representations of unanalysed speech when speech perception is challenging. Using DCM, Leff et al. (Leff et al., 2008) found that feedforward, rather than feedback, connections were associated with the difference between intelligible speech in quiet and unintelligible (time-reversed) speech. In addition, these lower-level regions have previously been associated with speech intelligibility in some studies: Binder et al. (2004) found that an anteriolateral temporal region—corresponding to Te1.1— correlated with accuracy on a phoneme-in-noise discrimination task, and Wild et al. (2012) found that activity in Te1.0 covaried with the intelligibility of degraded speech. The current results support the idea that lower-level processes (in Te1.0 and Te1.1) are associated with the perceptual challenge of a less favourable TMR during speech-in-noise perception. In addition, we show that perceptual challenges during figure-ground perception affect these lower-level processes in a similar way as do challenges during speech-in-noise perception.

A previous fMRI study of figure-ground perception (Teki et al., 2011), which used the same maps of auditory cortex that we used here, found no evidence for activity in primary auditory cortex— although they did not use a task, whereas we used a relevant, active task. A previous EEG study (O’Sullivan et al., 2015) found greater activity during active than passive listening to figure-ground stimuli, and a MEG study (Molloy et al., 2019) similarly found greater activity in primary auditory cortex under low than high visual load. Thus, our results are consistent with the idea that task effects modulate early auditory cortex during figure-ground perception. Here, we extend this idea by showing that the earliest stages of the auditory cortical hierarchy are more engaged (i.e., less inhibited) when the figure-ground task is more challenging due to a lower TMR.

Studies using other simultaneous or sequential stream segregation to study perceptual organisation have found that activity in both primary (Bidet-Caulet et al., 2007; Deike et al., 2010; Fishman et al., 2001; Micheyl et al., 2005; Schadwinkel and Gutschalk, 2010; Wilson et al., 2007) and non-primary (Deike et al., 2010; Schadwinkel and Gutschalk, 2010; Wilson et al., 2007) auditory cortex differs depending on how listeners perceive acoustic sources—for example the number of sources they perceive or which features of the scene they attend to. Also, Overath et al. (2010) found both primary and non-primary parts of auditory cortex were active when participants detected changes in spectrotemporal coherence in dense acoustic ‘textures’, which contain multiple components that changed frequency; participants made decisions about the how coherent the direction of frequency changes were across components.

The strongest—and highest probability—modulations of effective connectivity by difficulty were located in the left hemisphere. It is widely accepted that speech is processed bilaterally in auditory cortex (see Peelle, 2012; Scott and McGettigan, 2013). However, the left hemisphere modulations we observed are consistent with previous studies that have localised speech-in-noise effects to the left hemisphere. For example, Scott et al. (2004) found an area within the left anterior STG that showed a positive correlation with speech intelligibility, and Davis et al. (2011) found that the left posterior STG correlated positively with TMR. Regarding more basic stimuli designed to assess auditory scene analysis, two previous studies of sequential stream segregation found correlates of different percepts in the left but not right auditory cortex (Deike et al., 2010, 2004). Flinker et al. (2019) suggest that perceiving temporal modulations leads to left lateralised responses, whereas perceiving spectral modulations leads to right lateralised responses. Unlike other figure-ground tasks (Elhilali et al., 2009b; Gutschalk et al., 2008; Kidd, 1994; Kidd et al., 1995), the stochastic figure-ground task used here cannot be detected based on simple spectral separation; instead, it has been associated with a temporal coherence mechanism (Shamma et al., 2011; Teki et al., 2013). That we found predominantly left-hemisphere modulations therefore aligns both with the division proposed by Flinker et al. (2019), and with the findings from previous speech-in-noise tasks. However, it is worth noting that one right hemisphere intrinsic connection (Te1.1) was present in our DCM (Table 3), albeit with a lower probability (.71)—suggesting the effects are not entirely lateralised. One previous EEG study (Bidelman and Howell, 2016) reported greater right-hemisphere lateralisation when speech-in-noise was presented at a lower TMR, although in this study, participants were instructed to ignore the speech stimuli—and, therefore, responses are unlikely to relate to poorer intelligibility and may instead relate to stimulus acoustics.

In the current study, we manipulated difficulty by manipulating TMR, which is a naturally relevant quantity that varies greatly among different everyday listening settings. Rather than specifying a TMR that was fixed across participants, we selected TMRs for each participant that corresponded to 60% and 90% behavioural thresholds. This aspect of the design makes it less likely that the results reflect acoustic properties of different TMRs, but rather the perceptual challenges imposed by a lower TMR—which occur at different TMRs for different people. The selected TMRs and the acoustic noise (babble for speech-in-noise; random tone chords for figure-ground) also differed between the two tasks. Furthermore, absolute TMRs in each participant were regressed out of the model. Therefore, disinhibition of left Te1.0 and Te1.1 likely arose due to the increased difficulty associated with lower TMRs, rather than acoustic properties of the speech-in-noise and figure-ground tasks that covary with TMR.

Possibly, between-subjects differences in the disinhibition of left Te1.0 and 1.1 might help to explain why Holmes and Griffiths (2019) found that people who are worse at figure-ground perception are also worse at speech-in-noise perception. A common clinical observation is that patients report difficulties with speech-in-noise perception, despite no evidence of impaired peripheral function (Cooper and Gates, 1991; Hind et al., 2011; Kumar et al., 2007). Clinical measures are usually restricted to peripheral measures, such as the pure-tone audiogram, which do not fully account for individual differences understanding speech in noise (Cooper and Gates, 1991; Hind et al., 2011; Holmes and Griffiths, 2019; Kumar et al., 2007). Figure-ground perception has great potential as a useful clinical measure to predict speech-in-noise perception: Holmes and Griffiths (2019) demonstrate that it explains variance in speech-in-noise perception after accounting for differences in audiometric thresholds. One of the reasons it might be helpful in predicting speech-in-noise perception is because it assesses cortical processing that is relevant to speech-in-noise perception, which is not assessed by current clinical measures. In other words, some of the individual variability in speech-in-noise perception may arise from differences in disinhibition at the early stages of auditory cortical processing.

The results of the univariate analysis indicate that figure-ground perception predominantly activates a sub-set of the regions involved in speech-in-noise perception: both tasks reliably activate the superior temporal lobe (Table 2), but speech-in-noise perception leads to greater activity in bilateral STG, the left precentral gyrus, and the right cerebellum (Table 1). This finding is consistent with the idea that our figure-ground task is a basic version of speech-in-noise perception that relies on similar acoustic analysis (e.g., fundamental grouping processes), but does not require linguistic and articulatory processes involved in speech-in-noise perception.

The areas that were more strongly activated by speech-in-noise than figure-ground—bilateral STG, the left precentral gyrus, and the right cerebellum—have been associated with speech-in-noise tasks in previous studies. For example, the STG correlates positively with speech intelligibility (Scott et al., 2004) and TMR (Davis et al., 2011). Cerebellar activity has also been reported in previous studies (see Ackermann et al., 2007 for a review), despite the fact that—traditionally—the cerebellum is not commonly thought to be part of the speech network. Using PET, Salvi et al. (2002) found activity in the right cerebellum when speech-in-babble was compared to speech-in-quiet; perhaps, crucially, they selected levels for the speech and babble to ensure that performance was approximately 50% for each participant. Similarly, here, we ensured that speech intelligibility was below ceiling (60% or 90%) and equated performance between the speech-in-noise and figure-ground tasks. The involvement of the motor cortex (precentral gyrus) in speech perception has been long debated (see Scott et al., 2014), but several studies have found motor cortex activity during speech perception (e.g., Wilson et al., 2004; Wilson and Iacoboni, 2006). The current results lead further support to the claim that, compared to more basic auditory stimuli, speech perception is associated with activity in motor cortex.

Only two voxels showed greater activity for figure-ground than speech-in-noise perception, and they did not survive a stringent correction for family-wise error at an alpha of .001. However, it is worth noting that they are located close to parts of the IPS that have previously been associated with figure-ground perception (Teki et al., 2016, 2011). Consistent with these results, earlier work has demonstrated that IPS plays a role in basic auditory streaming (Cusack, 2005). Previously, IPS activity has been attributed to top-down attention (Cusack, 2005) or to perceptual ‘pop-out’ (Shamma and Micheyl, 2010; Teki et al., 2011) during auditory scene analysis. During figure-ground perception, predictions about frequencies can be very precise after the figure has been detected (because the frequencies of the figure remain the same for the entire figure duration), whereas speech changes frequency over time; thus, greater activity in IPS during figure-ground perception could reflect greater ‘pop-out’ of figures that remain the same frequency over time, than of speech, which changes frequency over time.

In summary, our results demonstrate common processes for figure-ground and speech-in-noise perception in early auditory cortex. We found that figure-ground perception predominantly activates a sub-set of regions involved in speech-in-noise perception. Modelling of BOLD responses showed that greater difficulty in both tasks is associated with disinhibition in left Te1.0 and left Te1.1—implying that the early stages of the auditory cortical hierarchy increase their gain when speech-in-noise and figure-ground perception become more difficult. Ultimately, these results suggest a common cortical substrate that links perception of basic and natural sounds—and might explain why people who are worse at figure-ground perception are also worse at speech-in-noise perception.

## Materials and Methods

### Subjects

49 participants completed the experiment. We measured their pure-tone audiometric thresholds at octave frequencies between 0.25 and 8 kHz in accordance with BS EN ISO 8253-1 (British Society of Audiology, 2004). We excluded one participant who had a mild sloping hearing loss; all other participants had 6-frequency average thresholds better than 20 dB HL in either ear. We also excluded 4 participants who were not native English speakers, leaving 44 participants—which is the number we aimed to analyse based on an *a priori* power analysis. A sample size of 44 was estimated using NeuroPower (http://neuropowertools.org) with power = 0.8, and was based on publicly available fMRI data reported by Hakonen et al. (2017). The 44 participants (23 male) we tested were 19–35 years old (median = 22.7 years; interquartile range = 6.0) and all reported that they were right-handed.

The study was approved by the University College London Research Ethics Committee, and was performed in accordance with relevant guidelines and regulations. Informed consent was obtained from all participants.

### Stimuli

Stochastic figure-ground stimuli were based on Holmes & Griffiths (2019). They contained 50-ms chords, gated by a 10-ms raised-cosine ramp, with 0 ms inter-chord interval. Each chord contained multiple pure tones at frequencies selected from a logarithmic scale between 179 and 7246 Hz (1/24th octave separation). The stimuli contained a figure that lasted 2100 ms (42 chords) and a background that lasted 3100 ms (62 chords). The background comprised 5–15 pure tones, whose frequencies were selected randomly at each time window. The figure comprised 3 components; the frequencies were selected randomly on each trial and were the same for the entire figure duration. The figure began 500 ms (10 chords) after the background. For half of stimuli, 4 chords (lasting 200 ms) were omitted from the figure. For the other half, the same number of components (3) were omitted from the background (4 chords; 200 ms). The omitted components began 20–42 chords after the onset of the figure-ground stimulus (10–32 chords after the onset of the figure); the components were always omitted while the figure was present, even if they were omitted from the background.

Sentences for the speech-in-noise task were from the English version of the Oldenburg matrix set (HörTech, 2014) and were recorded by a male native-English speaker with a British accent. The sentences are of the form “<Name> <verb> <number> <adjective> <noun>” and contain 10 options for each word (see Table 4). An example is “Rachel brought four large chairs”. Recorded sentences were normalised to the same root-mean-square amplitude and lasted on average 2.2 seconds (standard deviation = .1). The sentences were presented simultaneously with 16-talker babble, which began 500 ms before the sentence began, ended 500 ms after the sentence ended, and was gated by a 10-ms raised-cosine ramp. The babble was taken from a continuous track lasting 20 seconds; a different segment of the babble was selected on each trial.

**Table 4.**
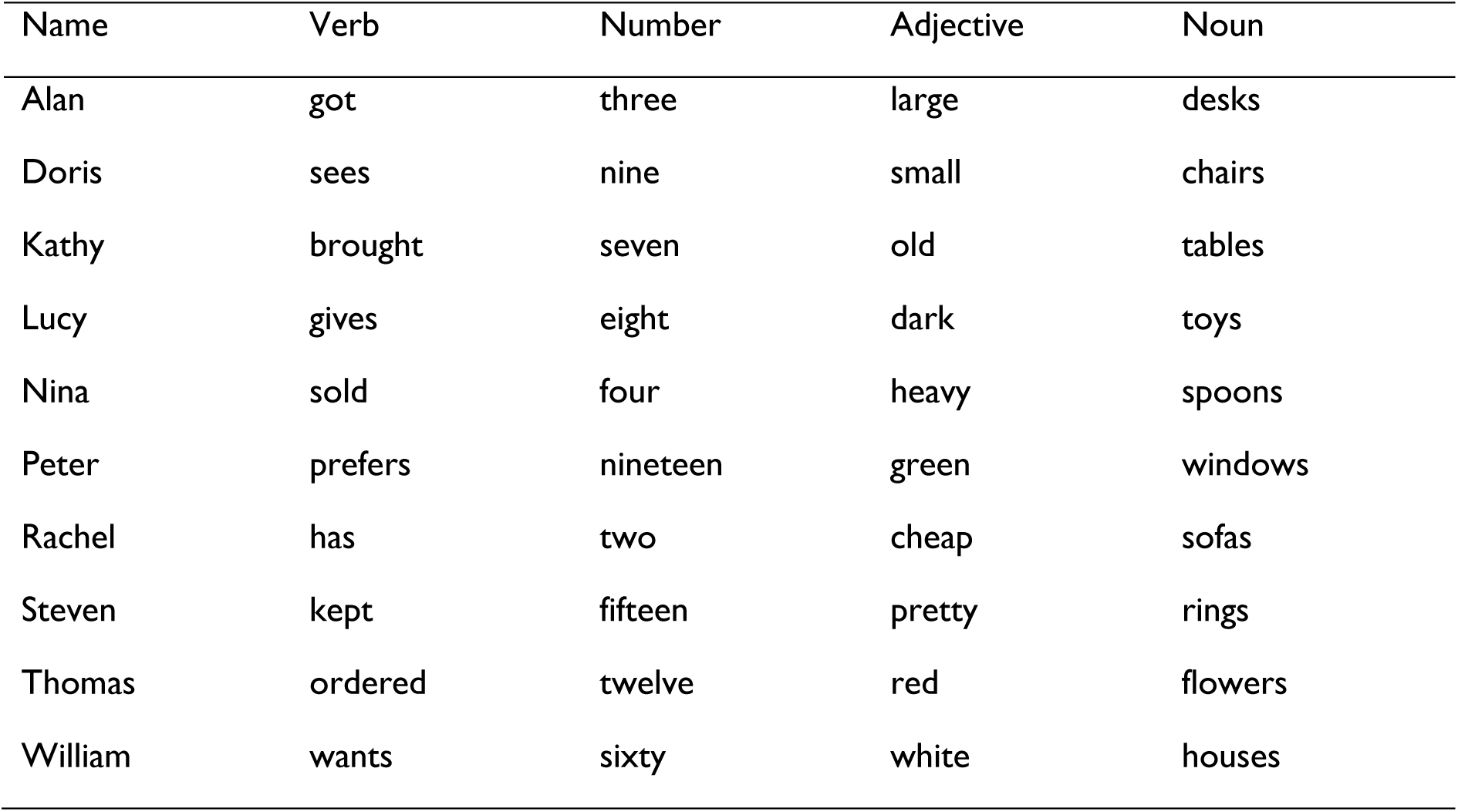
Words from the English version of the Oldenburg International Matrix corpus, which was used in the speech-in-noise task. Sentences were constructed by selecting one word from each of the five columns with equal probability, ensuring that transition probabilities between words were equated across sentences. Sentences were recorded in their entirety rather than as individual words.

| Name | Verb | Number | Adjective | Noun |
| --- | --- | --- | --- | --- |
| Alan | got | three | large | desks |
| Doris | sees | nine | small | chairs |
| Kathy | brought | seven | old | tables |
| Lucy | gives | eight | dark | toys |
| Nina | sold | four | heavy | spoons |
| Peter | prefers | nineteen | green | windows |
| Rachel | has | two | cheap | sofas |
| Steven | kept | fifteen | pretty | rings |
| Thomas | ordered | twelve | red | flowers |
| William | wants | sixty | white | houses |

Stimuli were presented using MATLAB (R2015a) and Psychtoolbox (version 3.0.14). Sounds were presented at 75 dB A, which was measured using a Brüel & Kjær (Nærum, Denmark) Type 2636 sound level meter. The levels were calibrated separately with the equipment that was used for the behavioural and MRI sessions.

### Experimental Procedures

#### Pre-scan behavioural

At the beginning of the experiment, participants completed a behavioural session, to determine their thresholds for 60% and 90% performance on the stochastic figure-ground (SFG) and speech-in-noise (SPIN) tasks.

The pre-scan behavioural was conducted in a sound-attenuating booth. Participants sat in a comfortable chair facing an LCD visual display unit (Dell Inc.). Acoustic stimuli were presented through a Roland Edirol UA-4FX (Roland Corporation, Shizuoka,, Japan) USB soundcard connected to circumaural headphones (Sennheiser HD 380 Pro; Sennheiser electronic GmbH & Co. KG).

Participants first performed a short (< 5 minute) block to familiarise them with the figure-ground stimuli. During the familiarisation block, they heard the figure and ground parts individually and together, with and without a gap in the figure.

After familiarisation, we determined thresholds for the two tasks. We varied the target-to-masker ratio (TMR) between the target (figure or speech) and masker (background tones or babble noise, respectively) in a weighted adaptive procedure (Kaernbach, 1991). We used a step size ratio of 6:1 to estimate 60% thresholds and a step size ratio of 9:1 to estimate 90% thresholds. We used four separate blocks to estimate the TMRs corresponding to 60% and 90% thresholds for the two tasks. Each block included two separate but interleaved runs, which were identical except that different stimuli were presented. Each run started at a TMR of 0 dB and terminated after 10 reversals. The step size began at 1 dB and decreased to .5 dB after 3 reversals. Identical stimuli were used in the 60% and 90% blocks, but they were presented in different orders.

To estimate figure-ground thresholds, participants completed a yes-no task. On each trial, participants heard a figure-ground stimulus and had to decide whether or not there was a gap in the figure. The figure contained a gap on 50% of trials. On trials in which there was no gap in the figure, there was a gap in the background of the same magnitude (for details, see ‘Stimuli’ section above). Participants responded by clicking buttons on the screen corresponding to yes and no responses.

During the speech-in-noise blocks, participants also completed a yes-no task. They had to decide whether a sentence written on the screen was the same as the target sentence they heard spoken. The written sentence was presented on the screen after the spoken sentence had ended, and was identical to the spoken sentence on 50% of trials. It was different on 50% of trials: On these trials, one word in the written sentence differed from the word in the spoken sentence; this word occurred at each position in the sentence with equal probability, and was selected randomly from the other words in the corpus. Participants responded by clicking buttons on the screen corresponding to yes (same sentence) and no (different sentence) responses.

The order of the figure-ground and speech-in-noise blocks were counterbalanced across participants. Before the first block of each task, participants performed a 6-trial practice at 3 dB TMR, with feedback.

#### MRI

The MRI session was completed on the same day, immediately after the pre-scan behavioural. The same figure-ground and speech-in-noise tasks were presented, but at fixed TMRs—corresponding to the adapted TMRs from the pre-scan behavioural. Each task was presented at two different TMRs: One corresponding to the 90% threshold (SPIN-90 and SFG-90) and another corresponding to the 60% threshold (SPIN-60 and SFG-60).

Participants laid on a bed in the MRI scanner. Visual stimuli were presented through an Epson EB-L1100U projector, which participants viewed through a mirror attached to the head coil. Auditory stimuli were presented through a Roland Edirol UA-4FX (Roland Corporation, Shizuoka,, Japan) USB soundcard connected to Ear-Tone Etymotic earphones (Etymotic Research, Inc., Illinois, U.S.A.) with disposable foam ear tips.

We presented 8 functional runs, each containing 24 trials. Each run contained 6 trials from each condition, which were pseudorandomly interleaved. Figure 4 shows a schematic of the trial structure. Each trial lasted 8 seconds and contained three major components: A visual cue, which indicated the task for the upcoming trial; the acoustic stimuli; and a probe sentence, which cued participants to make a response. We used a sparse sampling method with one functional MRI volume acquisition at the end of each trial. The acoustic stimuli were presented in the silent gap between scans. They began, on average, 1.4 seconds after the start of the trial and were jittered within an interval of 2 seconds (i.e., .4–2.4 seconds after the end of the previous scan). The visual cue was the word “figure” or “speech”; it was presented for .4 seconds (.35 seconds during the previous scan, and .05 seconds after the previous scan had ended). A fixation cross then appeared on the screen until the probe sentence was presented 5.6 seconds after the trial began. The probe sentence remained on the screen until the visual cue for the next trial began. On figure-ground trials, the probe sentence was “Gap in figure?”. On speech-in-noise trials, the probe was a written sentence followed by a question mark. Participants responded using a button box in their right hand; they pressed one button to respond “yes”—if the there was a gap in the figure (figure-ground task) or if the written sentence matched the spoken sentence (speech-in-noise task)—and a different button to respond “no”.

**Figure 4.**
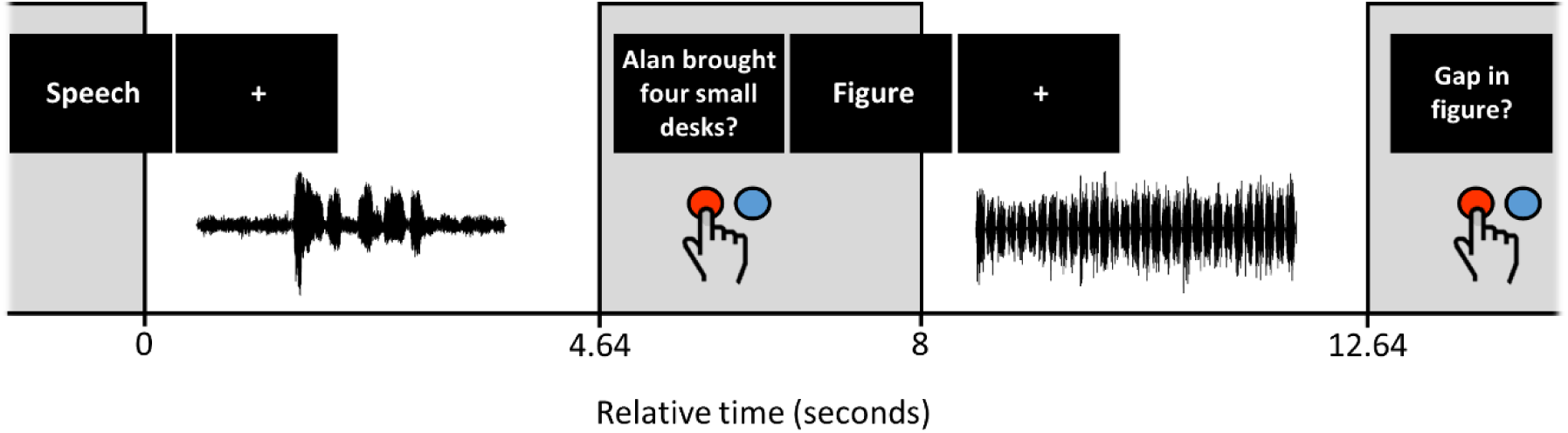
Schematic of the trial structure during the MRI session. The upper row displays the onset of visual stimulus presentation. The lower row displays the positioning of acoustic stimuli, and the approximate timing of button press responses. Grey bars represent functional volume acquisitions, which each lasted 3.36 seconds. Two exemplar trials are shown: a speech-in-noise trial followed by a figure-ground trial.

Before participants began the MRI session, they first completed a practice of 24 trials outside the scanner using the same equipment that was used for the pre-scan behavioural. The trial structure of the practice was identical to the MRI session, and participants received no feedback about their responses.

### MRI data acquisition

MRI was conducted on a 3.0 Tesla Siemens MAGNETOM TIM Trio MR scanner (Siemens Healthcare, Erlangen, Germany) at the Wellcome Centre for Human Neuroimaging (London, U.K.) with a 64-channel receive coil.

T2*-weighted functional images were acquired using echo-planar imaging (EPI) with field of view of 192 × 192 × 144 mm; voxel size = 3.0 × 3.0 × 2.5 mm; echo spacing of 30 ms; time-to-repeat (TR) of 3.36 seconds; 48 slices; anterior-to-posterior phase encoding; bandwidth of 2298 Hz/Px. Acquisition was whole-brain transverse oblique, angled away from the eyes. We used sparse temporal sampling (Hall et al., 1999): A delay of 4.64 seconds was imposed between successive volumes, such that each volume acquisition began 8 seconds after the previous volume acquisition began. We collected 24 volumes from each participant in each of the 8 runs, plus an additional ‘dummy’ scan, which was presented immediately prior to the first trial of each run and was excluded from the analyses. We collected field maps immediately after the functional runs (short TE = 10.00 ms and long TE = 12.46 ms).

At the end of the session, we acquired a whole-brain T1-weighted anatomical image (MPRAGE, 176 slices; voxel size = 1 mm isotropic; field of view 256 × 256 × 176 mm; PAT GRAPPA of factor 2; anterior-to-posterior phase encoding; TR = 2530 ms, TE = 3.34 ms).

### Analyses

#### Pre-scan behavioural

We calculated thresholds as the median of the last 6 reversals in each run. We averaged the thresholds from the two interleaved runs within each block.

#### Behaviour during scan

To determine behaviour during the MRI session, we calculated d′ (Green and Swets, 1966) with loglinear correction (Hautus, 1995). Trials with no response were included in the analysis as misses and false alarms (i.e., as incorrect responses).

#### MRI pre-processing

MRI data were processed using SPM12 (Wellcome Centre for Human Neuroimaging, London, UK). Each participant’s functional images (EPIs) were unwarped using their field maps and were realigned to the first image of the run. We then applied slice time correction. The functional and anatomical images were co-registered to the mean EPI, then normalised to the standard spm12 template (avg305T1). We spatially smoothed the images using a Gaussian kernel with 4 mm full-width at half-maximum.

#### Statistical parametric mapping

We modelled the fMRI timeseries for each participant with a General Linear (convolution) Model, with the motion realignment parameters as covariates of no interest. Each stimulus was modelled as a delta function, convolved with the canonical hemodynamic response function. We ran a contrast of all trials over the (implicit) baseline, which we used to define regions of interest (ROIs) for the DCM analysis. For completeness, we also ran contrasts to test the main effect of Task (figure-ground and speech-in-noise), the main effect of Difficulty (TMRs for 90% or 60% performance), and the interactions between Task and Difficulty. We entered the contrasts into a second level analysis, which applied one-sample t-tests at the group level.

#### Dynamic Causal Modelling

DCM is used to infer effective connectivity—and how directed causal influences among neural populations are affected by experimental manipulations. In brief, DCM is based on a model of neural population dynamics, which is combined with a hemodynamic model. The technical details of DCM are explained in other papers (see Friston et al., 2017, 2003; Zeidman et al., 2019a, 2019b). We first inferred the effective connectivity parameters that best fit each participant’s fMRI data, then estimated the parameters and their uncertainty at the group level using parametric empirical Bayes. Finally, we used a particular form of Bayesian model comparison—namely, Bayesian model reduction—to establish which model of effective connectivity best explain the group data.

Here, we were interested in making group-level inferences about how greater difficulty in figure-ground and speech-in-noise tasks modulates intrinsic (i.e., within-region) and extrinsic (i.e., between-region) connectivity (i.e., a main effect of Difficulty) and whether there are modulations specific to greater difficulty in one task over the other (i.e., an interaction between Task and Difficulty).

#### Selection of timeseries

We extracted timeseries for each subject in 8 ROIs (left and right Te1.0, Te1.1, Te1.2, and Te3). To ensure the timeseries showed reliable task-related activity—and did not include voxels with random signal fluctuations—we selected voxels for each subject that were significant at pre-specified thresholds at both the group and individual-subject levels. In detail, we masked the group-level contrast (All Trials > baseline) with anatomical masks extracted from the SPM Anatomy Toolbox (version 2.2c) (Eickhoff et al., 2005), corresponding to the 8 ROIs. We used the voxels that were below the *p* = .05 threshold (after family-wise error correction) to generate a functional mask for each ROI. We applied these functional masks to the individual-subject results, and retained voxels at the individual-subject level that were below a threshold of *p* = .05 uncorrected. Where no voxels within an ROI survived the p < .05 threshold (right Te1.0: 2/44 participants; right Te1.1: 11/44 participants; right Te1.2: 1/44 participants; left Te1.1: 2/44 participants; left TE1.2: 1/44 participants), we increased the individual-subject threshold in increments of .05 until one or more voxels survived. (Note that the thresholds applied in the selection of timeseries only specify the voxels that are included in the analysis and do not determine statistical significance of the DCM analysis.) Finally, we created a summary timeseries for each ROI in each participant by extracting the principal eigenvariate.

#### DCM estimation

The DCM for each participant included 8 nodes corresponding to the extracted timeseries for the 8 ROIs. We specified the input timing for the DCM analysis as vectors specifying All Trials of interest, the main effects of Task and Difficulty, the interaction between Task and Difficulty, and the motion and run covariates. We estimated (using Bayesian model inversion with Variational Laplace) a fully-connected DCM for each participant, which included all possible combinations of intrinsic and extrinsic fixed connections. We allowed All Trials and the main effect of Task to serve as external (i.e., direct or driving) inputs to each node. We allowed the main effect of Difficulty and the interaction between Task and Difficulty to modulate all intrinsic and extrinsic connections, and serve as external inputs to each region.

#### Group level inference

To estimate parameters at the group level (i.e., across participants), we took the parameters of interest for each participant to a second-level Parametric Empirical Bayes (PEB) analysis. This is a hierarchical model of connectivity parameters, with connectivity parameters from all subjects at the first-level and a GLM at the second-level, estimated using a variational scheme. Our first-level parameters of interest were the modulations of intrinsic and extrinsic connectivity by the main effect of Difficulty and by the Task–Difficulty interaction (i.e., parameter matrix B from each subject’s DCM). We entered the absolute TMRs in each of the four conditions for each participant (which differed according to their thresholds measured in the pre-scan behavioural session) as regressors in the second level of the PEB model. Having estimated parameters of the full PEB model, we then pruned away parameters using Bayesian Model Reduction (BMR)—which performs an automatic (‘greedy’) search over the model space, essentially comparing the evidence for reduced models that have particular parameters ‘switched off’. The model evidence considers both accuracy (how well the model fits the data) and complexity (the difference between model parameters and their prior values, which were always set to zero). The algorithm iteratively discards parameters if the reduced model has greater evidence. Thus, simpler models (i.e., those with more parameters ‘switched off’) that fit the data sufficiently accurately are preferred, because greater complexity reduces the model evidence. The final 256 models from the BMR were entered into a Bayesian Model Average (Hoeting et al., 1999; Penny et al., 2006), which performs a weighted average of parameters across models, according to the posterior probabilities (Pp) of the models. We discounted parameters whose posterior probabilities were less than .95, to focus our conclusions on high-probability parameters; although, for completeness, we report the probabilities of all parameters in Table 3.

## Acknowledgements

This work was supported by WT091681MA and DC000242-31 to Timothy D. Griffiths. This work was conducted at the Wellcome Centre for Human Neuroimaging, which is funded by Wellcome (Ref: 203147/Z/16/Z). The funders had no role in study design, data collection and analysis, decision to publish or preparation of the manuscript.

## Author Contributions

E.H., P.Z., K.J.F, and T.D.G. designed the study, interpreted the data, and wrote the manuscript. E.H. collected and analysed the data.

